# Unprecedented incidence of *Trypanosoma cruzi* infections in a cohort of dogs directly detected through longitudinal tracking at multi-dog kennels, Texas, USA

**DOI:** 10.1101/2021.06.24.449798

**Authors:** Rachel E. Busselman, Alyssa C. Meyers, Italo B. Zecca, Lisa D. Auckland, Andres H. Castro, Rebecca E. Dowd, Rachel Curtis-Robles, Carolyn L. Hodo, Ashley B. Saunders, Sarah A. Hamer

## Abstract

Canine Chagas disease, caused by the protozoan parasite *Trypanosoma cruzi*, is increasingly recognized as a health concern for dogs in the USA, and infected dogs may signal geographic regions of risk for human disease. Dogs living in multi-dog kennel environments where triatomine vectors are endemic may be at high risk for infection. We monitored a cohort of 64 *T. cruzi*-infected and uninfected dogs from across 10 kennels in Texas, USA, to characterize changes in infection status over time. We used robust diagnostic criteria in which reactivity on multiple independent platforms was required to be considered positive. Among the 30 dogs enrolled as serologically- and/or PCR-positive, all but one dog showed sustained positive *T. cruzi* diagnostic results over time. Among the 34 dogs enrolled as serologically- and PCR-negative, 10 new *T. cruzi* infections were recorded over a 12-month period. The resulting incidence rate was 30.7 *T. cruzi* infections per 100 dogs per year. This study highlights the risk of *T*. *cruzi* infection to dogs in kennel environments, despite multiple vector control methods employed by kennel owners. To protect both dog and human health, there is an urgent need to develop more integrated vector control methods as well as prophylactic and curative antiparasitic treatment options for *T. cruzi* infection in dogs.

## Introduction

The protozoan parasite *Trypanosoma cruzi*, agent of Chagas disease, is a vector-borne zoonotic pathogen endemic to the Americas. The pathogen infects an estimated eight million people and hundreds of mammal species (1). *T. cruzi* is predominantly transmitted in the feces of infected triatomines (‘kissing bugs’) through contact with wounds or mucus membranes or ingestion of infected insects or fecal material (1). Oral transmission is thought to be the most important route in domestic dogs and wild mammals and is a highly efficient mode of transmission (2–5).

Enzootic cycles of *T. cruzi* transmission have been documented in the southern USA, where opossums (*Didelphis virginiana*), woodrats (*Neotoma* spp.), coyotes (*Canis latrans*), and other mammals naturally infected with *T. cruzi* come into contact with endemic triatomine species (6–8) and maintain parasite transmission (1, 9–12). The wildlife species involved in the transmission of *T. cruzi* often inhabit peridomestic environments and share spaces with domestic dogs outdoors, increasing dogs’ risk of exposure to infected vectors (10, 13). In settings where dogs share spaces with humans in and around homes, dogs serve as sentinels for human infection, maintaining *T. cruzi* transmission and amplifying disease in domestic and peridomestic environments (14, 15). Thus, areas of high domestic dog seroprevalence may indicate areas of heightened risk for human infection (16).

In the USA, *T. cruzi* infection has been reported in dogs across the southern states (17–19). Studies from Texas, Oklahoma, and Louisiana report dog infection prevalence from 3.6-22.1%, and up to 57.6% in some multi-dog kennels (19–27). Infection with *T. cruzi* shows no strong breed predilection, and shelter and stray dogs are likely exposed to *T. cruzi* often due to their high level of exposure to vectors, resulting in high prevalence of infection (19, 22, 28, 29). Additionally, government working dogs along the USA-Mexico border are exposed to *T. cruzi* with a seroprevalence of up to 18.9% (26), in which infections have led to fatal cardiac disease (30). In both humans and dogs, Chagas disease causes a range of clinical symptoms, progressing through acute and chronic stages of disease, which can include severe heart disease and death (3, 21, 31). While insect vectors are endemic to the southern USA, canine travel introduces a veterinary health concern as infected dogs move outside of endemic areas to areas where clinicians are less familiar with the symptoms of Chagas disease (32).

While measures of infection prevalence from cross-sectional studies are useful in quantifying the burden of disease in populations, the incidence rate (i.e. the number of new infections per population per unit time) can provide a direct measurement of risk. Direct measurements of incidence in natural animal populations are rare, as they require tracking of healthy, uninfected individuals over time to monitor for new infections. Alternatively, incidence can be inferred indirectly by comparing seroprevalence across different age cohorts of animals. Using this indirect approach based on serology, we are aware of two studies investigating the *T. cruzi* incidence rate in dog populations in the USA. The first estimated a serologic incidence of 3.8% in juvenile dogs along the Texas-Mexico border (33); the second estimated an incidence rate of 2.3 new cases per year in dogs of all ages in Louisiana shelters (19). We used a longitudinal study design to directly measure incidence of *T. cruzi* infection in dogs of Central/South Texas, an area with a high risk of infection based on suitable triatomine habitat and autochthonous human cases (34). Uncertainty about transmission risk and the lack of an effective treatment or vaccine pose significant problems to veterinarians and dog owners following diagnoses of Chagas disease. In this study, we tracked the serostatus and PCR-status of a matched cohort of *T. cruzi*-positive and negative dogs at three intervals over 12-months, allowing for the direct detection of *T. cruzi* infection in dogs in kennel environments.

## Methods

### Locations and Sample Collection

All samples were obtained from privately-owned animals in accordance with guidelines approved by the Texas A&M University’s Institutional Animal Care and Use Committee and Clinical Research Review Committee (2018-0460 CA). Using a prospective cohort study design, we enrolled a network of 10 multi-dog kennels throughout Central and South Texas with a prior history of *T. cruzi* infection in at least one of their dogs. Kennel locations were categorized by Texas ecoregions (28). Dogs at these kennels were bred and trained primarily to aid hunting parties or compete in American Kennel Club dog events and were classified according to American Kennel Club breed groups. Enrollment criteria included dogs of any breed or sex at least one year of age, in residence at one of the selected kennels, and negative for other selected vector-borne infectious diseases (see below).

We aimed to identify and enroll approximately two to four *T. cruzi*-infected and two to four *T. cruzi*-uninfected dogs at each kennel for longitudinal tracking at approximate 6-month intervals. During the initial sampling time point between May and July 2018, we tested 134 dogs (between seven and 20 dogs at each of the 10 kennels) for evidence of *T. cruzi* and four other vector-borne infections. Based on the enrollment criteria, 64 dogs were identified as meeting the criteria and were enrolled in the study. When possible, *T. cruzi*-positive and *T. cruzi*-negative dogs were frequency (group) matched on the basis of age, sex, and breed across all kennels. Each blood sampling event consisted of collecting approximately 5mL of blood in an EDTA (ethylenediaminetetraacetic acid) tube and 4mL in a clotting tube. The second sampling event occurred between December 2018 and March 2019, and the third sampling event occurred between May and September 2019. All 10 kennels were visited at each time point to collect blood samples from enrolled dogs.

### Serology

At each of the three time points, serum samples were tested for *T. cruzi* antibodies using three serological tests: Chagas Stat-Pak (ChemBio, Medford, NY, USA), Chagas *Detect Plus* Rapid Test (InBios International, Inc., Seattle, WA, USA), and an indirect fluorescent antibody (IFA) test. The Chagas Stat-Pak and Chagas *Detect Plus* are rapid tests that, while not labeled for use in dogs, have been shown to be sensitive and specific to detect *T. cruzi* antibodies and are commonly used in dogs for research purposes (25, 35, 36). Serum samples were tested according the manufacturer’s instructions, and results were determined after 15 minutes. In all cases, the integrated ‘control’ bands appeared as expected on the rapid tests. Rapid tests were considered negative when the ‘test’ band had no color development or when a very faint band- not perceptible enough to be a clear positive- developed. The IFA was performed by the Texas A&M Veterinary Medical Diagnostic Laboratory (TVMDL) using 200 μL of serum to test for IgG antibodies against *T. cruzi*. Reactivity on at least two of the three serologic tests was one of the criteria (in addition to PCR positivity, see below) to enroll or consider a dog *T. cruzi*-positive. An IDEXX 4Dx test (IDEXX Laboratories, Inc., Westbrook, ME, USA) was run to exclude any dogs that may have had other vector-borne infections, including *Anaplasma* spp., *Erlichia* spp., *Borrelia burgdorferi*, and *Dirofilaria immitis*.

### DNA Extraction, Quantification, and PCR

A DNA extraction procedure from large volumes (5mL) of EDTA-treated blood was used in an effort to increase sensitivity of detection of *T. cruzi* using the E.Z.N.A Blood DNA Maxi kit (Omega Bio-Tek, Norcross, GA, USA) according to the manufacturer’s protocol, except 650μL of elution buffer was used. *T. cruzi-n*egative controls (phosphate buffered saline) were included in the DNA extractions. For each extracted sample, DNA was quantified using a spectrophotometer (Epoch, BioTek Instruments, Inc., Winooski, VT, USA). In the case of low DNA yield from the MAXI extraction (<85 ng/μL), or if contamination was suspected in the PCR, a second extraction was performed using 250μL of clotted blood using the E.N.Z.A. Tissue DNA kit, according to the manufacturer’s protocol, except 50 μL of elution buffer was used (Omega Bio-Tek).

Samples were tested using qPCR for the presence of *T. cruzi* satellite DNA using the Cruzi 1/2 primer set and Cruzi 3 probe in a real-time assay, which amplifies a 166-bp segment of a repetitive nuclear DNA (37, 38) as previously described (26). This PCR method is sensitive and specific for *T*. *cruzi* when compared to other PCR techniques (39). *T. cruzi* DNA extracted from isolate Sylvio X10 CL4 (ATCC 50800, American Type Culture Collection [ATCC], Manassas, VA, USA; DTU TcI) was used as the *T. cruzi*-positive control in all PCRs. The qPCR amplification curves were checked for quality (sigmoidal curves), and a sample was considered *T. cruzi*-positive if the Ct value was lower than 36 (40).

### Statistical Analyses

Dogs were categorized as *T. cruzi*-positive at a given time point if they met the criteria for serological positivity (positive on at least two of three independent serologic assays) and/or if they met the criteria for PCR positivity (Ct value lower than 36). We calculated the Kappa index to determine the agreement among all three serological tests (19, 26). To describe the risk of *T. cruzi* infection in the population of dogs that were initially enrolled as *T. cruzi*-negative, we calculated the incidence rate as the number of new *T. cruzi* infections per 100 dogs per year. To reflect the time *T. cruzi*-negative dogs in the population were at risk, 0.5 years was subtracted from the population at risk (n = 34) for each of the three *T. cruzi*-negative dogs lost to follow up after the 6-month time point (n = 3).

## Results

### Dog Enrollment

Of the 134 dogs screened, we enrolled 30 *T. cruzi*-positive and 34 *T. cruzi*-negative dogs across 10 kennels in Central and South Texas (Table 1). Kennels were located in one of four ecoregions, primarily (7/10) in the South Texas Plains. Dogs enrolled in the study were between the ages of 13 months and 12.2 years (mean = 6.1 years; median = 6.1 years). Dogs were frequency matched based on age, sex, and breed: age was matched within 8 months (average *T. cruzi*-positive = 6.6 years; average *T. cruzi*-negative = 5.8 years), sex was matched within four dogs (positive = 14 males and 16 females; negative = 19 females and 15 males), and breeds were matched between *T. cruzi*-positive and *T. cruzi*-negative dogs (various hound breeds were considered “Hound” for matching purposes; three positive and negative Belgian Malinois, two positive and negative German Shorthair Pointers, three positive and negative Hounds, one positive and seven negative Labrador Retrievers, two positive and negative Brittany Spaniels, and 19 positive and 17 negative English Pointers per group). During the initial screening of 134 dogs across all 10 kennels, one dog was positive for *Dirofilaria immitis* and was excluded from the study. In total, five dogs were lost to follow up after the 6-month time point: two dogs enrolled as *T. cruzi*-negative (including one that seroconverted to positive by 6-months) moved locations and were sold to new owners, and three dogs (one *T. cruzi*-positive and two negative) had died.

**Table 1:**
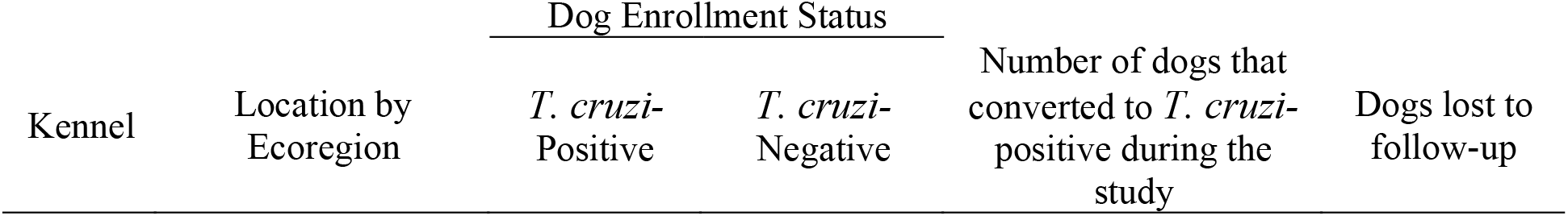
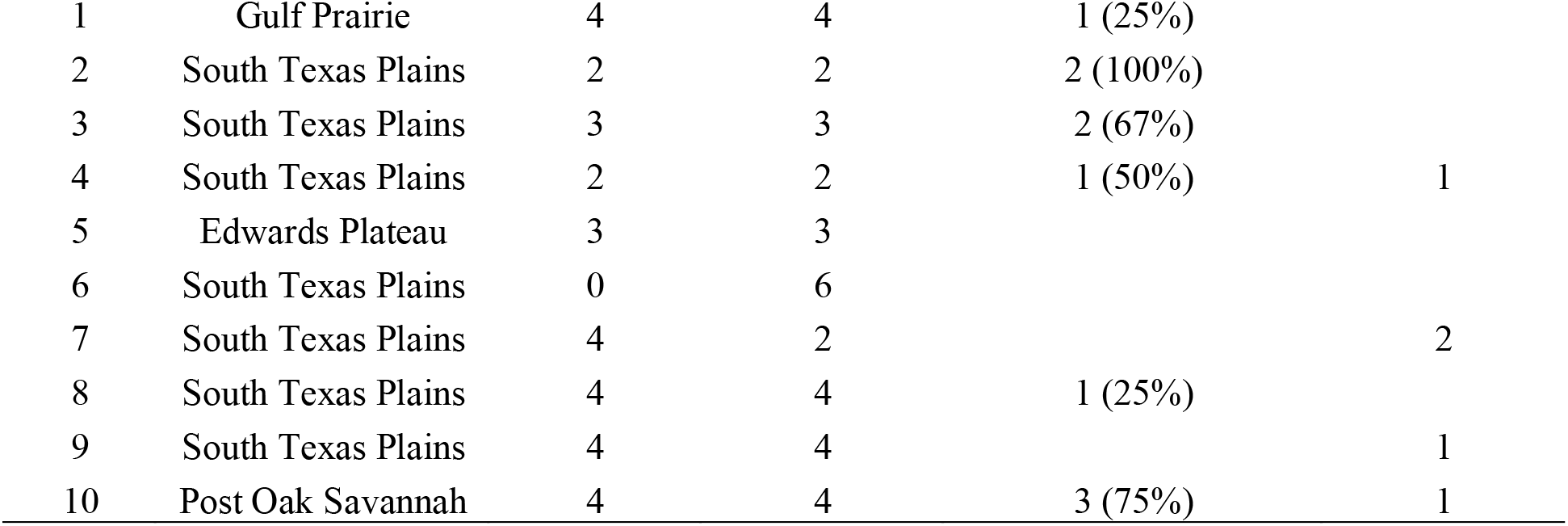
Location of enrolled dogs and number of conversion events from each kennel.

Eluted DNA concentrations from MAXI extractions used in the study ranged from 88.6 ng/μL to 348.5 ng/μL (n = 104; mean = 172.4 ng/μL; median = 164.2 ng/μL).

Of the 30 dogs enrolled as *T. cruzi*-positive at the initial time point, 26 (86.7%) were positive on all three serologic tests; two (6.7%) were positive on both rapid tests yet negative on IFA; and one (3.3%) was positive on a single rapid test and had a high titer (>1280) on IFA (Table 2). Additionally, one dog (D76) was enrolled as *T. cruzi*-positive based solely on PCR positivity (3.3%; Table 2). Ten of the aforementioned *T. cruzi*-seropositive dogs were also PCR positive. The overall prevalence of PCR-positivity at the time of enrollment among dogs enrolled as *T. cruzi*-positive was 11/30, 36.7% (Table 3).

**Table 2:** Serology of the 30 dogs enrolled as *T. cruzi*-positive in the study. The two rapid tests (Chagas Stat-Pak and Chagas *Detect Plus*) are scored as Positive (“+”) or Negative (“−”), and the IFA titer is reported in which a result of <20 is noted as Negative (“−”).

| Dog ID | Initial Time Point |  |  | 6-month Time Point |  |  | 12-month Time Point |  |  |
| --- | --- | --- | --- | --- | --- | --- | --- | --- | --- |
|  | Chagas Stat-Pak | IFA Titer | Chagas *Detect Plus* | Chagas Stat-Pak | IFA Titer | Chagas *Detect Plus* | Chagas Stat-Pak | IFA Titer | Chagas *Detect Plus* |
| D05 | + | 5120 | + | + | 640 | + | + | 1280 | + |
| D06 | + | 5120 | + | + | 2560 | + | + | 1280 | + |
| D07 | + | 2560 | + | + | 160 | + | + | 640 | + |
| D08 | + | 2560 | + | + | 640 | + | + | 2560 | + |
| D17 | + | - | + | + | - | + | + | - | + |
| D21 | + | 5120 | + | + | 2560 | + | + | 640 | + |
| D26 | + | 320 | + | + | 160 | + | + | 160 | + |
| D27 | + | 2560 | + | + | 640 | + | + | 5120 | + |
| D30 | + | - | + | + | - | + | + | - | + |
| D39 | + | 160 | + | + | - | + | + | 20 | + |
| D42 | + | 640 | + | + | 160 | + | + | 640 | + |
| D43 | + | 640 | + | + | 320 | + | + | 320 | + |
| D49 | + | 640 | + | + | ≥1280 | + | + | 320 | + |
| D54 | + | 80 | + | + | 160 | + | + | 80 | + |
| D75 | + | 640 | + | + | 320 | + | + | 5120 | + |
| D76* | - | - | - | - | - | - | - | - | - |
| D77 | + | 1280 | + | + | 320 | + | + | 5120 | + |
| D79 | + | 2560 | + | + | 320 | + | + | 5120 | + |
| D94 | + | 1280 | + | + | 80 | + | + | 640 | + |
| D96 | + | 2560 | + | + | 640 | + | + | 5120 | + |
| D99 | + | 1280 | + | + | 320 | + | + | 320 | + |
| D100 | + | 640 | + | + | 160 | + | + | 320 | + |
| D103 | + | 1280 | + | + | 160 | + | + | 640 | + |
| D104 | + | 1280 | + | + | 640 | + | + | 2560 | + |
| D108 | + | 2560 | + | + | 640 | + | + | 5120 | + |
| D110 | + | 320 | + | + | 160 | + | + | 320 | + |
| D121 | + | 320 | + | + | 320 | + | + | 160 | + |
| D122 | + | 320 | + | + | 160 | + | + | 80 | + |
| D123 | + | 80 | + | + | 40 | + | + | 160 | + |
| D124 | - | >1280 | + | - | 640 | + | Deceased |  |  |

\*D76 was serologically-negative but enrolled as *T. cruzi*-positive due to PCR positivity in the blood.

**Table 3:**
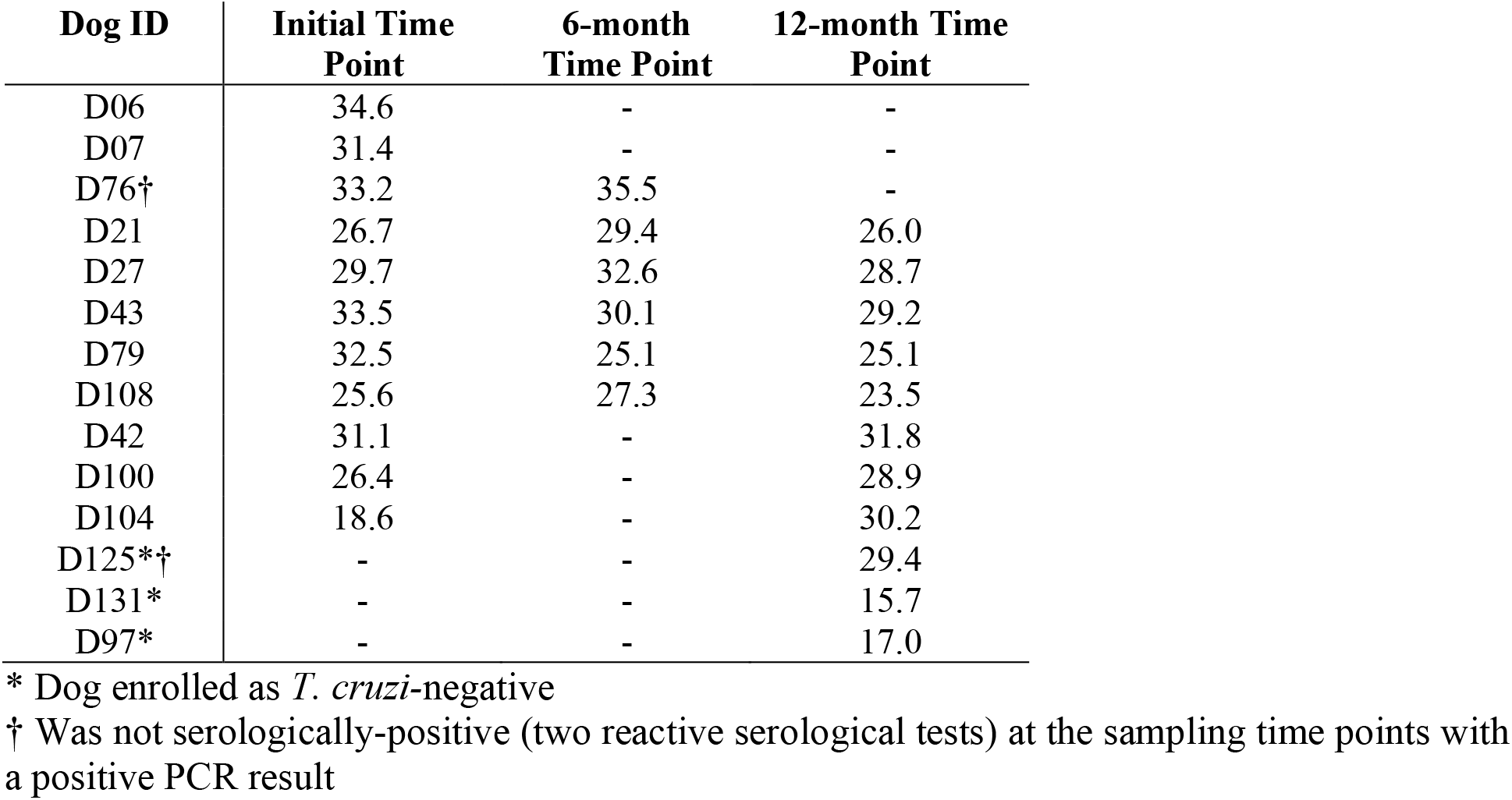
PCR results for dogs with a Ct value indicating a *T. cruzi*-positive sample. No Ct value or a Ct value of 36 or higher during qPCR is considered a negative PCR result and is noted as “−” in this table.

Of the 34 dogs enrolled as *T. cruzi*-negative, 31 (91.2%) were negative on all three serologic tests and PCR at the initial time point. Three dogs (3/34, 8.8%) were weakly positive on a single rapid test (two dogs were positive on only the Chagas Stat-Pak, and one dog was positive only on the Chagas *Detect Plus*).

### Longitudinal Tracking

Twenty-nine of the 30 dogs enrolled as *T. cruzi*-positive maintained their *T. cruzi*-positive status at the 6-month and 12-month time points across the study with minor fluctuations in individual serologic and PCR test results over time; for example, one dog (D39) was positive on all three serological assays during the initial and 12-month time point but positive on only the two rapid tests during the 6-month time point (Tables 2 and 3). The dog enrolled as *T. cruzi*-positive based on PCR-positivity only (D76) was again PCR-positive at the 6-month time point but PCR-negative at the 12-month time point and never developed antibodies detectable on any of the three serologic assays. Of the 11 dogs enrolled as *T. cruzi*-positive that had PCR-positive results at the initial time point, five dogs (45.4%) were consistently PCR-positive at all three time points, 3 (27.3%) were PCR-positive at only the first and third sampling trips, one dog (9.1%) was PCR-positive at only the first and second visits, and two dogs (18.2%) were only PCR-positive at the initial time point (Table 3).

Of the 34 dogs enrolled as *T. cruzi*-negative, 24 (70.6%) remained negative at the 6-month and 12-month time points (Figure 1). One dog (2.9%) remained serologically-negative but PCR-positive at the 12-month time point, and nine dogs (26.5%) were seropositive by the end of the study (Figure 1). Notably, two dogs (5.9%) were seropositive on both rapid serologic assays at the 6-month time point and therefore met the definition for seroconversion, but were seropositive on only one rapid serologic assay by the 12-month time point. In total, three dogs enrolled as *T. cruzi*-negative (8.8%) were PCR-positive at the 12-month time point (Figure 1).

**Figure 1:**
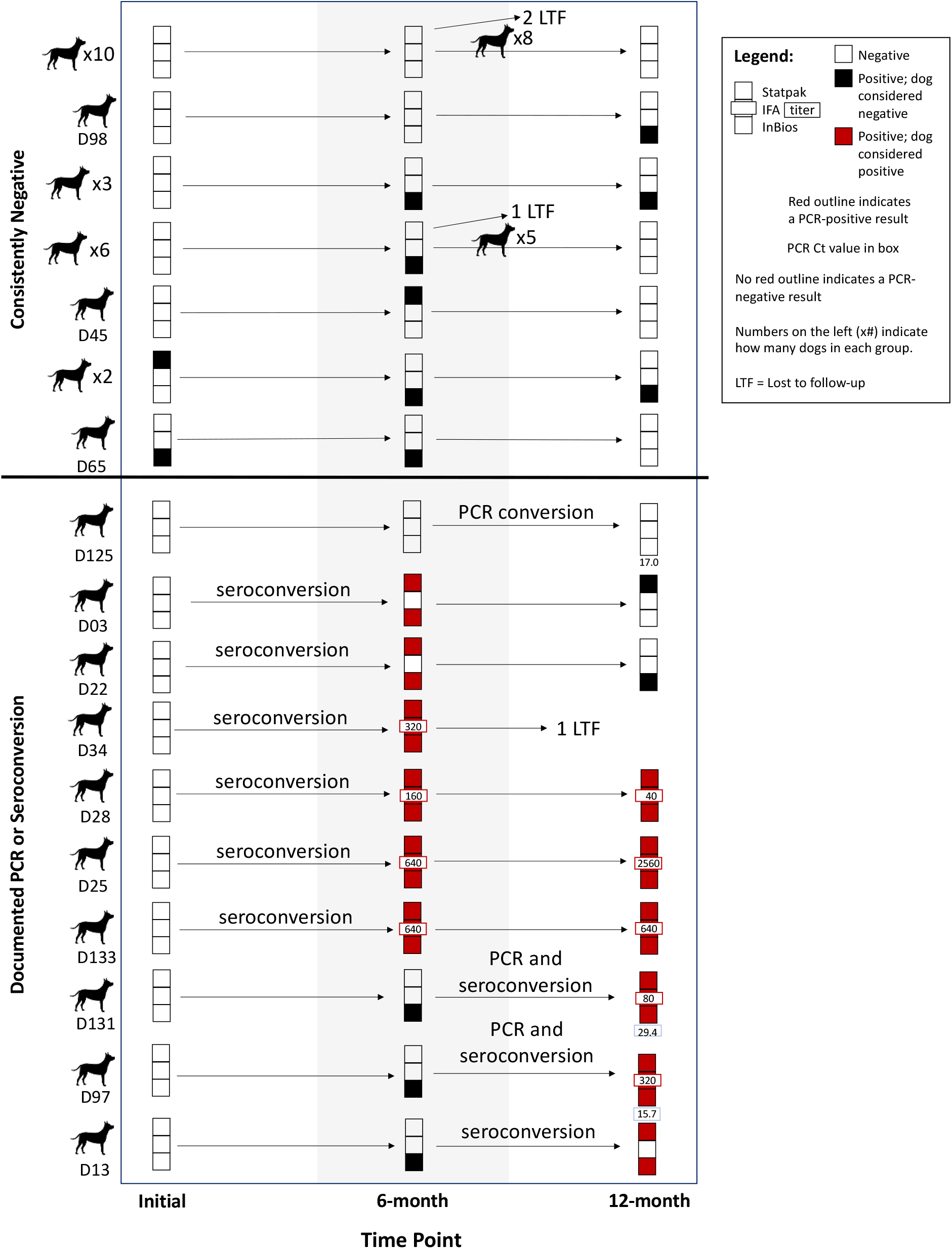
Serology and PCR results of all dogs enrolled as *T. cruzi*-negative (n = 34) at each sampling time point. When multiple dogs had the same progression of test results, this progression is depicted as a single row in the figure with the number of dogs shown on the left side of the figure. Changes in serostatus or PCR status are written between time points in which the conversion took place. The titer for positive IFA tests and the Ct value for PCR-positive tests are noted in the symbol. Dog icon is in the public domain.

The Kappa Indices comparing serological tests across all time points, showed the Chagas Stat-Pak and IFA had almost perfect agreement (kappa = 0.84), while there was substantial agreement between the Chagas *Detect Plus* and IFA, and Chagas *Detect Plus* and Chagas Stat-Pak (kappa = 0.65 and 0.70 respectively).

### Incidence

A total of 10 out of the 34 (29.4%) dogs enrolled as *T. cruzi*-negative converted to positive throughout the one-year study. After adjusting the time the total population was at risk to account for attrition of three *T. cruzi*-negative dogs after the 6-month time point, this results in an incidence rate of 30.7 *T. cruzi* infections per 100 dogs per year. If the two dogs that seroconverted by the 6-month time point but only maintained positivity on a single test at the 12-month time point (D03 and D22) were not true conversions, then the incidence rate would be 24.6 new *T. cruzi* infections per 100 dogs per year.

## Discussion

We characterized the *T. cruzi* serologic and PCR-status of 64 dogs at 3 time points over the course of one year and demonstrated a high incidence of *T. cruzi* infection in a cohort of dogs housed in multi-dog kennels across Texas. In our cohort, 34 dogs were negative at enrollment, and we recorded 10 new infections (29.4%) resulting in an incidence rate of 30.7 new infections per 100 dogs per year. Otherwise stated, a dog at these kennels has a risk of 30.7% of acquiring *T. cruzi* infection within one year.

Direct quantification of *T. cruzi* incidence in dogs is rare, likely due to the challenges of collecting longitudinal data (41). More commonly, incidence is estimated indirectly. For example, researchers conducted a serosurvey of 540 dogs in Louisiana and found a 6.9% prevalence of *T. cruzi* infection (19). Assuming each animals’ exposure to *T. cruzi* infection is similar between animals and over time, they also estimated the incidence of new *T. cruzi* infection in dogs was 2.3 new infections per 100 dogs/year. This estimate is much lower than our calculated incidence but similar to the estimated incidence in juvenile dogs along the Texas-Mexico border (33). In that study, the *T. cruzi* prevalence in juvenile dogs under 6 months old was used to estimate a *T. cruzi* serologic incidence of 3.8%, assuming all infections that occurred in dogs under 6 months old were newly acquired infections. This methodology does not consider congenital infections, and further, can only be applied to the juvenile age group. In contrast to the previous studies estimating incidence, our longitudinal study allowed us to directly detect new *T. cruzi* infections using serology and PCR over one year, and thus report a direct measure of incidence in a closed population of dogs.

There is no gold standard to test for *T. cruzi* infections in dogs, and the IFA is the only test approved for veterinary Chagas diagnostics. While the Chagas Stat-Pak and the Chagas *Detect Plus* are often used for research purposes, discordant test results are common in canine testing for Chagas disease (26). As recommended by the WHO for human Chagas diagnostics (42), reactivity on multiple serologic assays was the criteria used in our study to consider a dog *T. cruzi*-positive. We observed substantial to high agreement across serologic tests, reflecting the strict inclusion criteria and matched study design, where *T. cruzi*-positive dogs were enrolled as positive only when there was some agreement across serologic assays. Overall, the Chagas Stat-Pak and IFA had the best agreement with an overall kappa index of 0.84 (Table 4); whereas, there was reduced pairwise agreement between the other combinations of tests (kappa range 0.65-0.70). Cross-sectional studies of dog *T. cruzi* serostatus have found less agreement, even high discordance, between Chagas tests using the Kappa index, with only slight to moderate agreement between tests (19, 26). Given the absence of gold standard antemortem testing methods, our data suggest a multi-test approach is still necessary in dog Chagas disease diagnostics. PCR was used in addition to serologic tests as a diagnostic tool for *T. cruzi* in this study, with the ability to detect new infections prior to antibody formation. The PCR protocol used in this study was shown to be one of the most sensitive and specific PCR methods to detect *T. cruzi* based on an international survey of PCR methods of *T. cruzi* detection (39) and has been previously used to detect *T. cruzi* in blood samples (38). We expect to see fewer PCR positives than antibody positive results because PCR-positivity indicates active circulation of the parasite in the animals’ blood, which may occur for only short time periods in infected dogs (3). By using four independent tests to categorize animals as *T. cruzi*-positive or negative, and repeating sampling at 3 time points to look for change in infection status, our study design was robust and sensitive to capture new infections in a way that has not before been done in veterinary Chagas studies in the United States.

**Table 4:** Kappa Index table to statistically show pairwise comparisons and agreement among three serological tests: Chagas Stat-Pak, Chagas *Detect Plus*, and indirect fluorescent antibody (IFA) test.

|  |  |  |  |
| --- | --- | --- | --- |
| A. | IFA+ | IFA - | Total |
| Chagas Stat-Pak + | 86 | 13 | 99 |
| Chagas Stat-Pak - | 2 | 86 | 88 |
| Total | 88 | 99 | 187 |
| Almost perfect agreement: | **0.84** |  |  |
| B. | IFA+ | IFA - |  |
| Chagas *Detect Plus* + | 88 | 33 | 121 |
| Chagas *Detect Plus* - | 0 | 66 | 66 |
| Total | 88 | 99 | 187 |
| Substantial agreement: | **0.65** |  |  |
| C. | Chagas Stat-Pak + | Chagas Stat-Pak - |  |
| Chagas *Detect Plus* + | 96 | 25 | 121 |
| Chagas *Detect Plus* - | 3 | 63 | 66 |
| Total | 99 | 88 | 187 |
| Substantial agreement: | **0.70** |  |  |

In this study, one dog (D76) was PCR-positive yet antibody negative at enrollment, indicating the presence of *T. cruzi* DNA in the blood at the time of sampling, and thus this dog was enrolled as *T. cruzi*-positive. We expected this dog to develop detectable antibodies by the time of the 6- and 12-month time points. However, the dog did not seroconvert and displayed a weak PCR-positive result (high-positive Ct value) at the 6-month time point. The reasons for a lack of seroconversion in this dog are unknown but could be due to health conditions that resulted in a suppressed immune system or other health measures we did not assess as part of this study, or the initial and subsequent PCR-positive results were not reflective of a true infection.

Two dogs (D03 and D22) met the definition of incident cases because of their initial enrollment as *T. cruzi*-negative (each had one very faint band bands on one test; see Methods; classified as negative) and seroconversion at the 6-month time point, at which time they were reactive on both rapid serologic assays. However, at the 12-month time point, both dogs were only positive on one serologic assay (D03 on the Chagas Stat-Pak; D22 on the Chagas *Detect Plus*), and therefore failed to meet the diagnostic criteria for positivity at that time point, which required reactivity on at least two independent tests. As a new infection followed by self-cure would be unlikely (1, 41), we suspect this scenario reflects the limitations of using unvalidated diagnostic tests for dogs, and future prospective sampling from these dogs would be enlightening. If these two dogs represent false conversions, this means our incidence rate would be lower, at 24.6 new infections per 100 dogs/year. Even an incidence only considering eight new infections (out of 34) is markedly higher than previously estimated dog incidence rates for *T. cruzi* infections. Additionally, the three dogs enrolled as negative, despite having a weakly positive band on a single rapid test, remained negative through the course of the study, with either one weakly positive test or no positive test at each timepoint, providing evidence in this study that a single weak band is not likely to signal an early infection or an infection that will become more diagnostically apparent the year to follow. The need for improved diagnostic tools for detection of *T. cruzi* infection is apparent for human medicine (43), and our study highlights the need in veterinary medicine as well (26).

Over the course of the study, five dogs were lost to follow up either due to death or to moving locations. Unfortunately, necropsies were not performed and postmortem samples were not available; thus, the causes of death (and potential role of Chagas disease) remain unknown. One of the deceased dogs was *T. cruzi*-positive, and the owners described a slow decline over two months with observed weight loss, but no veterinary visits or treatment were elected by the owners.

There was geographic variation in the frequency of incident cases across the 10 studied kennels. Kennels varied from no incident infections to three of the four dogs enrolled as *T. cruzi*-negative having newly acquired *T. cruzi* infection. The incident cases were distributed across six of the 10 kennels. This variation may indicate natural variation in triatomine abundance or infection among kennels (44), or the importance of other factors, such as vector control around the kennels, kennel integrity, or exposure to bugs while working and when outside of kennels.

There is no approved pharmaceutical treatment or prevention for Chagas disease in dogs, and diagnosis often occurs during the chronic phase when experimental treatments rarely change the outcome of disease and treatment is designed to manage symptoms (3). As no vaccines or approved treatment options exist for dogs, and cardiac symptoms can cause acute death or chronic illness (45), preventing contact with infected vectors is paramount; this requires a thorough understanding of transmission cycles and risk. Accordingly, Chagas disease prevention and control is largely focused on reducing a dog’s contact with vectors. Approaches to halting contact include vector control with insecticides applied to kennel outdoor spaces, keeping dogs indoors, avoiding outdoor lights that attract bugs, and altering vegetation to reduce the density of wildlife reservoirs and kissing bugs. However, various vector control measures are challenging, as no insecticides are specifically labeled for use in the control of triatomines in the USA. In particular, it is common for adult kissing bugs to be encountered in kennel environments, posing a transmission risk to the dogs (44). Triatomines have proven difficult to eradicate even in areas with robust vector control programs, as locating the bugs is difficult, time consuming, and sometimes unsuccessful (46, 47).

Multi-dog kennels should be areas of targeted intervention in the USA, as the incidence of dog infection is high. We found dogs at many of these kennels have a high risk of *T. cruzi* infection over the course of one year, highlighting the need to develop thorough vector control methods that can be deployed in kennel environments and identify, test, and make available prevention and treatment options for *T. cruzi* infection in dogs in the USA. The loss of hunting and working dogs has an economic impact on breeders, trainers, and ranch owners, and developing next steps to prevent or treat canine Chagas disease not only is necessary for veterinary health but to decrease the human impact of Chagas disease in dogs. Further, travel of infected dogs from endemic regions of the southern USA to non-endemic regions, where there is less awareness of Chagas disease, is occurring (32), in which case diagnosis may be harder to attain. Because dogs can achieve a level of parasitemia sufficient to infect kissing bugs during acute disease, there is concern for infected dogs serving as a local reservoir to perpetuate the parasite life cycle (22, 24, 48). Accordingly, any interventions to reduce infection in dogs and improve their overall health may also have the add-on benefit of reducing risk of locally acquired human disease as well.

## Acknowledgements

The American Kennel Club Canine Health Foundation Grant No 02448 provided funding. We thank participating kennel owners for allowing access to dogs. We also thank Carlos Rodriguez at the Texas Veterinary Medical and Diagnostic Lab for conducting the IFAs and Ester Carbajal for fieldwork assistance.

